# Identification of KKL-35 as a novel carnosine dipeptidase 2 (CNDP2) inhibitor by *in silico* screening

**DOI:** 10.1101/2025.09.27.674371

**Authors:** Takujiro Homma, Koki Shinbara, Tsukasa Osaki

**Affiliations:** Department of Pharmacology, Graduate School of Medicine, Osaka Metropolitan University, Osaka 545-8585, Japan; Bara Pharma, Tokyo 141-0031, Japan; Department of Biochemistry and Molecular Biology, Graduate School of Medical Science, Yamagata University, 2-2-2 Iidanishi, Yamagata 990-9585, Japan

**Author notes:** Correspondence author Takujiro Homma. These authors contributed equally to this work (Takujiro Homma, Koki Shinbara).

**Keywords:** CNDP2, glutathione, *in silico* screening

## Abstract

Extracellular glutathione (GSH) is degraded on the cell surface, in which the γ-glutamyl residue is removed to generate cysteine–glycine (Cys–Gly) dipeptides that are subsequently transported to the cytoplasm. Carnosine dipeptidase II (CNDP2) is a cytoplasmic enzyme that hydrolyzes Cys–Gly and plays an important role in maintaining intracellular cysteine (Cys) homeostasis. CNDP2-mediated hydrolysis of Cys–Gly promotes Cys mobilization and contributes to the replenishment of intracellular GSH levels. CNDP2 is frequently overexpressed in various cancers and has been implicated in tumor cell proliferation and progression. This mechanism may enhance cancer cell survival by causing resistance to oxidative stress, which indicates that CNDP2 is a potential therapeutic target for cancer treatment.

Although bestatin (BES) has been identified as a CNDP2 inhibitor, its limited specificity and suboptimal drug-like properties have limited its therapeutic potential. In this study, we performed an *in silico* screen of a small-molecule compound library and identified KKL-35 as a novel CNDP2-binding molecule. Molecular dynamics (MD) simulations suggested that KKL-35 interacts within the catalytic pocket. Biochemical assays confirmed that it inhibits CNDP2 enzymatic activity, albeit with lower potency compared with BES. Despite its modest intrinsic activity, KKL-35 exhibits favorable physicochemical and pharmacokinetic properties, which are characterized by a low topological polar surface area (TPSA), reduced molecular flexibility, and well-balanced lipophilicity. This positions it as an attractive and tractable starting point for lead optimization. Taken together, these findings establish KKL-35 as a validated CNDP2 inhibitor and a promising lead compound for the development of more selective therapeutics targeting CNDP2-mediated cancer cell metabolism.

## 1. Introduction

Carnosine dipeptidase 2 (CNDP2) is an intracellular enzyme that hydrolyzes cysteinyl-glycine (Cys-Gly) to release cysteine (Cys) [1], which is the rate-limiting amino acid required for the synthesis of the major intracellular antioxidant glutathione (GSH) [2]. GSH is degraded extracellularly by γ-glutamyl transferase (GGT), which removes the γ-glutamyl group [3]. Cys-Gly remains, which can be transported into cells and subsequently processed by CNDP2. This metabolic route plays an important role in maintaining intracellular redox homeostasis by sustaining cysteine availability for GSH resynthesis.

We previously reported that CNDP2 expression is upregulated in cells lacking xCT, which is the cystine/glutamate antiporter that imports oxidized cystine [4]. We also observed that, in a lack of Cys, cells compensate by hydrolyzing Cys–Gly to liberate Cys. Consistent with this mechanism, CNDP2-deficient cells exhibit heightened sensitivity to ferroptosis, which is an iron-dependent, lipid peroxidation–driven form of programmed cell death [5]. Notably, the potential role of CNDP2 as a regulator of oxidative stress has been demonstrated in mouse and Drosophila models [4,6]. Taken together, these results position CNDP2 as a key contributor to the cellular defense against oxidative stress.

Besides maintaining redox balance, recent studies have implicated CNDP2 in tumor biology. CNDP2 expression is increased in various cancers and is frequently associated with malignancy. For example, in ovarian cancer, CNDP2 promotes tumor growth and metastasis by activating the PI3K/AKT signaling pathway [7]. In colon cancer, CNDP2 expression upregulation facilitates cellular proliferation [8]. In laryngeal carcinoma, CNDP2 is considered a potential diagnostic and therapeutic marker [9]. Moreover, increased levels of CNDP2 protein have been detected in bronchoalveolar lavage fluid from patients with small-cell lung cancer, which further supports its relevance to malignancy [10]. Functional studies have indicated that CNDP2 knockdown inhibits tumor growth; however, the exact mechanisms underlying its oncogenic roles remain unclear.

Recent studies have revealed that CNDP2 facilitates tumor cell fitness through a cooperative nutrient scavenging mechanism, particularly under conditions of glutamine deprivation [11]. Because many cancers depend on glutamine for growth [12], tumor cells secrete CNDP2 to degrade extracellular oligopeptides into free glutamine. These metabolites act as public goods to sustain the proliferation of secreting and neighboring cancer cells, which is an important strategy for glutamine-addicted tumors. Genetic ablation or pharmacological inhibition of CNDP2 suppresses peptide-dependent growth *in vitro* and reduces tumor burden *in vivo*, whereas glutamine supplementation restores these phenotypes. This indicates that CNDP2 is a key mediator of nutrient recycling and metabolic adaptation.

Because of its dual roles in antioxidant defense and nutrient supply, CNDP2 has emerged as a promising therapeutic target; however, the availability of inhibitors, such as bestatin (BES; ubenimex), which is a *Streptomyces abikoensis* metabolite, is limited due to a lack of specificity. BES has been studied in acute myeloid leukemia and lymphedema and inhibits the degradation of various regulatory peptides, including oxytocin, vasopressin, and enkephalins. BES is a competitive, reversible inhibitor of multiple peptidases, including arginyl aminopeptidase, leukotriene A4 hydrolase, and membrane alanyl aminopeptidase, as well as membrane dipeptidase [13–15]; however, this broad target profile limits its use as a selective CNDP2 inhibitor.

Computational approaches are important to identify targeted compounds. Of these approaches, molecular docking combined with structure-based *in silico* screening provides a robust strategy for the identification of candidate compounds capable of interacting with the active sites of target proteins, thereby serving as a valuable approach in the early stages of drug discovery [16–19]. In this study, we combined virtual screening with a molecular dynamics simulation pipeline to identify novel CNDP2 inhibitors with improved specificity and therapeutic potential for cancer treatment. This *in silico* approach facilitates rational compound selection and provides a foundation for the development of next-generation CNDP2-targeted therapies.

## 2. Materials and Methods

### Protein Preparation for *In Silico* Screening

The target protein (CNDP2) was retrieved from the Protein Data Bank (PDB ID: 4RUH) and prepared for *in silico* screening using UCSF Chimera 1.18 [20]. Nonstandard molecules, including water and ions, except Mn, were removed because Mn is necessary for its activity [21]. Protein stability was evaluated by a Ramachandran plot using PDBsum [22]. Although molecular dynamic (MD) simulation was used to obtain a stable structure, the results did not differ remarkably. Therefore, the PDB structure with non-standard small molecules removed, except for Mn, was used for in silico screening.

### Library Preparation

The Selleckchem Bioactive Compound Library-I (Selleckchem, Houston, TX, USA) was screened. Most of the compounds in this library are commercially available, which enables rapid experimental validation through inhibition assays immediately after *in silico* screening and MD assessments. The library was filtered using Lipinski’s Rule of Five, which provides an empirical guideline for drug-likeness [23]. This rule states that orally active compounds typically have a molecular weight ≤ 500 Da, LogP ≤ 5, no more than 5 hydrogen bond donors, and no more than 10 hydrogen bond acceptors. Any molecule violating more than one of these criteria was removed. PAINS (Pan-Assay Interference Compounds) filters implemented in RDKit were also applied [24]. These filters identify and exclude molecules with substructures known to cause frequent false-positive results in biological assays, thereby improving the reliability of downstream screening. Compounds that passed both filters were subjected to structural preprocessing. Water molecules and inorganic ions associated with the compounds were removed, all hydrogens were added, and the Gasteiger partial charges were assigned to approximate the electronic distribution of each molecule for docking calculations [25]. Three-dimensional conformations were generated and subjected to geometry optimization using Merck Molecular Force Field 94 (MMFF94) [26]. MMFF94 is a general-purpose force field designed to reproduce experimental geometries and conformational energies across a broad range of organic molecules. This optimization step ensures that the ligands adopt physically reasonable low-energy conformations before the docking and MD simulations are conducted.

### *In Silico* Screening

The compound library was subjected to molecular docking using Smina, an optimized fork of AutoDock Vina that was designed for efficient virtual screening [27]. Docking was conducted within a cubic grid box (center coordinates: x = 18.82, y = 108.45, z = 39.05; box size: 15 Å). The grid box defines the search space where the ligand is allowed to explore possible binding poses. The grid box center was selected to coincide with the previously reported binding position of BES. Therefore, the screening conditions were designed to reproduce the reported docking pose of BES, which was included as a positive control. The exhaustiveness parameter was set to 8, which balances the thoroughness of search and computational efficiency by controlling the number of independent docking runs. Of the screened compounds, KKL-35 was identified as a binder with a docking pose comparable to that of BES. The docking results were analyzed and visualized. PyMOL (Schrödinger, LLC, 2015) was used to examine the three-dimensional binding conformations, whereas PoseEdit [28] generated two-dimensional interaction diagrams to facilitate the interpretation of hydrogen bonds, hydrophobic contacts, and other important ligand–protein interactions.

### MD Simulation and MM-PB/GBSA Free Energy Calculation

MD simulations were conducted using GROMACS 2024 to assess the stability of the protein–ligand complexes [29,30]. An ff14SB force field was applied to the protein, whereas the ligands were parameterized using ACPYPE based on the General AMBER Force Field (GAFF) with Gasteiger charges [31–33]. The complexes were embedded in a cubic water box with a 12 Å buffer, solvated with TIP3P water molecules, and neutralized with counter-ions [34]. Energy minimization was performed for 1,000 steps using the steepest descent algorithm, which iteratively reduces steric clashes by moving atoms along the negative gradient of potential energy. The systems were subjected to NVT equilibration for 20 ps at 310 K using Langevin dynamics, which is a stochastic integrator that maintains temperature stability by combining deterministic Newtonian motion with random forces and friction. Temperature coupling was controlled using the V-rescale thermostat to ensure accurate canonical ensemble sampling [35,36]. Subsequently, NPT equilibration was performed for 100 ps at 310 K and 1 bar with the C-rescale barostat, which is a stochastic velocity rescaling algorithm that provides robust sampling of the isothermal–isobaric ensemble [37]. Finally, MD simulations were conducted for 100 ns under periodic boundary conditions at 310 K and 1 bar, enabling unbiased observation of the protein–ligand interactions at the atomistic level. The trajectories were re-centered and corrected for periodic boundary artifacts, and the structural stability and binding affinity were assessed thoroughly by RMSD, RMSF, RoG, and hydrogen bond analyses, as well as MM-PBSA free energy and per-residue decomposition analyses using gmx_MMPBSA.

### *In Silico* Pharmacokinetic and Toxicological Analysis

BES and KKL-35 were selected and evaluated for their physicochemical properties, chemical features, and drug-likeness. Their Absorption, Distribution, Metabolism, Excretion, and Toxicity (ADMET) profiles were also assessed using the online servers, SwissADME and pkCSM [38,39].

### TargetPrediction

SwissTargetPrediction [40] was used to predict potential molecular targets of BES and KKL-35. The 2D structures of each compound were uploaded as query inputs, and predictions were conducted using the default parameters.

### Cell Culture and Chemicals

The HK-2 human proximal tubular cell line was obtained from the RIKEN Bioresource Center (Tsukuba, Japan). They were maintained in high-glucose DMEM (FUJIFILM Wako Pure Chemical, 044-29765) supplemented with 10% fetal bovine serum (FBS; Biowest, Riverside, MO, USA) and penicillin–streptomycin (FUJIFILM Wako Pure Chemical; 168-23191) at 37°C in a humidified incubator containing 5% CO_2_.

### Lentiviral Expression and FLAG Affinity Purification of CNDP2

The pENN.AAV.CB7.CI.CNDP2-flag.WPRE.rBG plasmid was obtained from Addgene (Plasmid #132685) and used as a template to amplify the mouse CNDP2-flag sequence. The amplified CNDP2-flag fragment was cloned into the pLVpro-EF1α lentiviral expression vector (Takara Bio) using In-Fusion Snap Assembly Master Mix (Takara Bio). Lentiviral particles encoding CNDP2-flag were generated by co-transfecting pLVpro-EF1α-CNDP2-flag with the pLVpro™ Packaging Mix (Takara Bio) into Lenti-293X cells, based on the manufacturer’s instructions. Following transfection, the culture medium was harvested and filtered to obtain a viral supernatant containing lentiviral particles. CNDP2-flag was overexpressed in HK-2 cells by lentiviral transduction. The expressed protein was purified from cell lysates using the DDDDK-tagged Protein Magnetic Purification Kit (MBL Life Science, Japan) based on the manufacturer’s instructions. The purified protein was analyzed by SDS-PAGE, and the presence of CNDP2 was determined by Coomassie Brilliant Blue (CBB) staining.

### Analysis of Dipeptidase Activity

The enzymatic activity of CNDP2 was measured in a 20-μL reaction mixture, which contained 0.1 M ammonium bicarbonate buffer (pH 7.0), 20 μg/mL purified CNDP2, and 10 mM dithiothreitol (DTT), along with either BES (Tokyo Chemical Industry, U0111; 0.0001–0.1 mM) or KKL-35 (Selleck, S6562; 0.1–1 mM). After preincubating for 10 min on ice, Cys–Gly (Cayman, #43262; 5 mM) and either MnCl_2_ (0.1 mM) or EDTA (0.1 mM) were added. The reactions were incubated at 37°C for 30 min and treated with 20 μL of 20 mM *N*-ethylmaleimide (NEM) in 50 mM ammonium bicarbonate for 10 min at room temperature, followed by the sequential addition of 40 μL methanol, 40 μL of the internal standard solution (10 μM L-methionine sulfone and 10 μM GSH-*N*-methylmaleimide in methanol), and 80 μL of chloroform. After vortexing and centrifuging at 12,000 × *g* for 15 min at 4°C, the upper aqueous phase was removed and dried in a SpeedVac concentrator.

The dried extracts were subsequently dissolved in deionized water and analyzed by liquid chromatography–tandem mass spectrometry (LC–MS/MS) using a Q-Exactive hybrid quadrupole–Orbitrap mass spectrometer (Thermo Fisher Scientific) equipped with a heated electrospray ionization source. The instrument was operated in the positive ionization mode. The LC system (Ultimate 3000, Dionex) consisted of a WPS-3000 TRS autosampler, a TCC-3000 RS column oven, and an HPG-3400RS quaternary pump. Separation was achieved using a SeQuant ZIC-pHILIC column (2.1 × 150 mm, 5 μm particle size; Merck KGaA). Mobile phase A was 20 mM ammonium bicarbonate (pH 9.8), and mobile phase B was 100% acetonitrile.

System control and data acquisition were done using Xcalibur 2.2 software. The raw data were processed using Compound Discoverer 2.1 (Thermo Fisher Scientific) and matched against the mzVault metabolite database (February 2017 version) to assess compound composition, based on accurate mass and isotopic patterns. Tentative metabolite identification was achieved by comparing the full MS and MS/MS fragment ions and validated using authentic standards. The compounds were grouped with a mass tolerance of 20 ppm and a retention time tolerance of 1 min. They were quantified based on their normalized peak areas. Glycine and Cys-NEM were quantified using calibration curves generated with standards and normalized to the internal standards.

## 3. Results

### Overview of the Screening Workflow

The strategy used for the discovery of CNDP2 inhibitors is presented in Fig. 1. The workflow consisted of sequential computational and experimental steps, beginning with molecular docking and its visualization in 2D and 3D, followed by MD simulations of the candidate compounds. Estimation of ligand-binding affinities by MM-PBSA and hotspot analysis was conducted to refine the list of putative inhibitors. The candidates were assessed for their pharmacokinetic and toxicity profiles using ADMET analysis, and enzyme inhibition assays were performed to measure their activity. This systematic approach ensured that only structurally compatible, pharmacologically favorable compounds progressed through each stage of the analysis.

**Figure 1.**
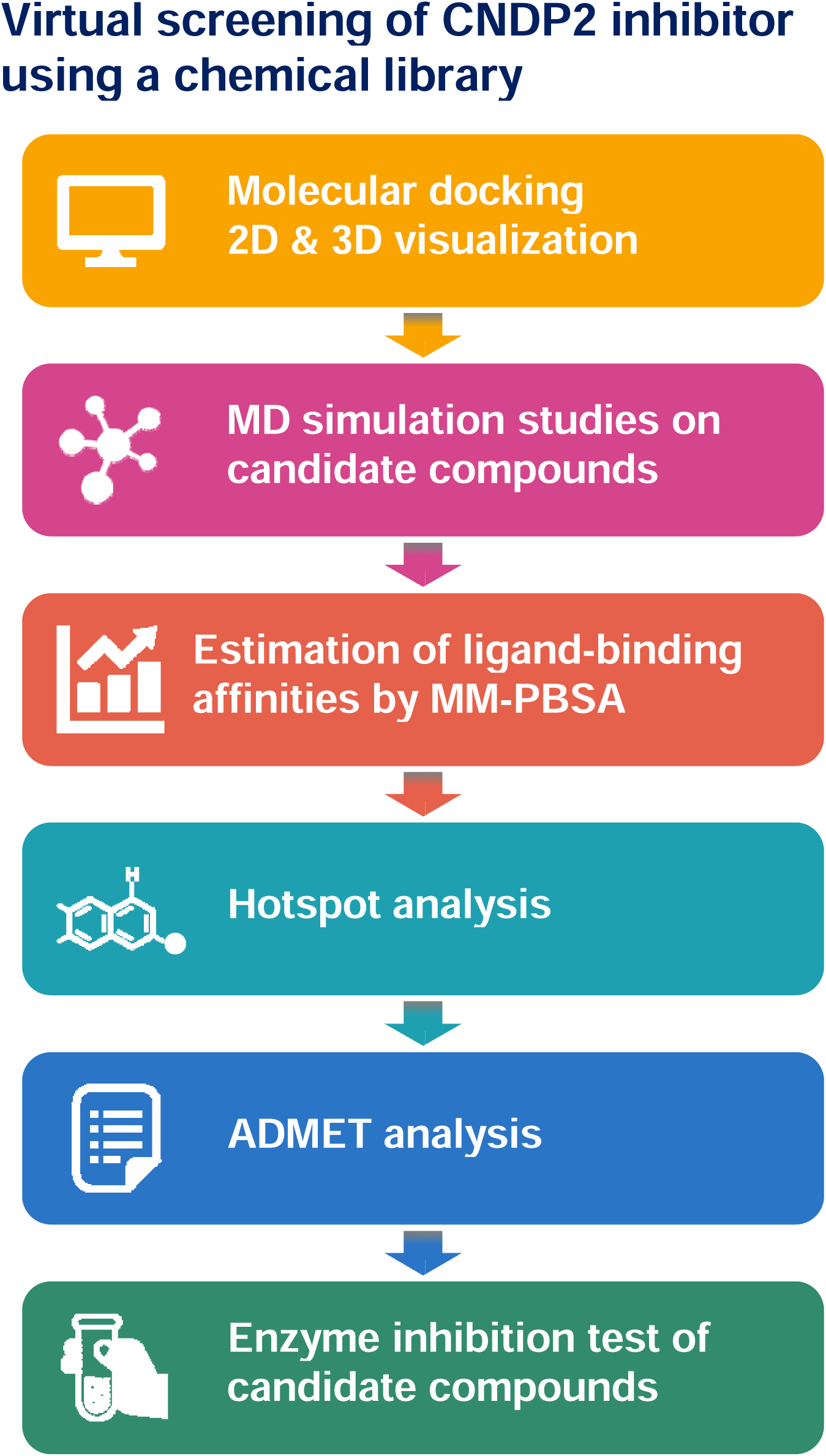
The discovery workflow for CNDP2 inhibitors, including structure validation, molecular docking, molecular dynamics (MD) simulations, binding free energy calculations, and ADMET analysis.

### Structural Validation of CNDP2

To implement this workflow, we validated the stereochemical quality of the CNDP2 enzyme structure (PDB: 4ruh) using PROCHECK analysis. The Ramachandran plot indicated that 91.4% of the non-glycine and non-proline residues were located in the most favored regions, with 8.2% and 0.3% in additionally and generously allowed regions, respectively. Only a single residue (0.1%) was observed in the disallowed region, which indicates minimal structural anomalies. Because a reliable protein model should have more than 90% of residues in the favored regions, the CNDP2 structure fulfilled this criterion [41]. G-factor analysis revealed an overall average of +0.11, further confirming the absence of unusual stereochemical parameters [42]. Taken together, these results validated CNDP2 as a high-quality structure appropriate for subsequent docking and MD simulations. The Ramachandran plot of the CNDP2 enzyme is shown in Fig. 2. The three-dimensional architecture of CNDP2 was also examined to visualize the docking environment. CNDP2 forms a dimeric structure with a well-defined catalytic cleft (Fig. 3A). BES are consistently localized within this catalytic pocket at the interface of the two domains. An enlarged view of the boxed region (Fig. 3B) highlights the close spatial arrangement of the ligands with key residues, thus supporting their compatibility with the inhibition of enzymatic activity.

**Figure 2.**
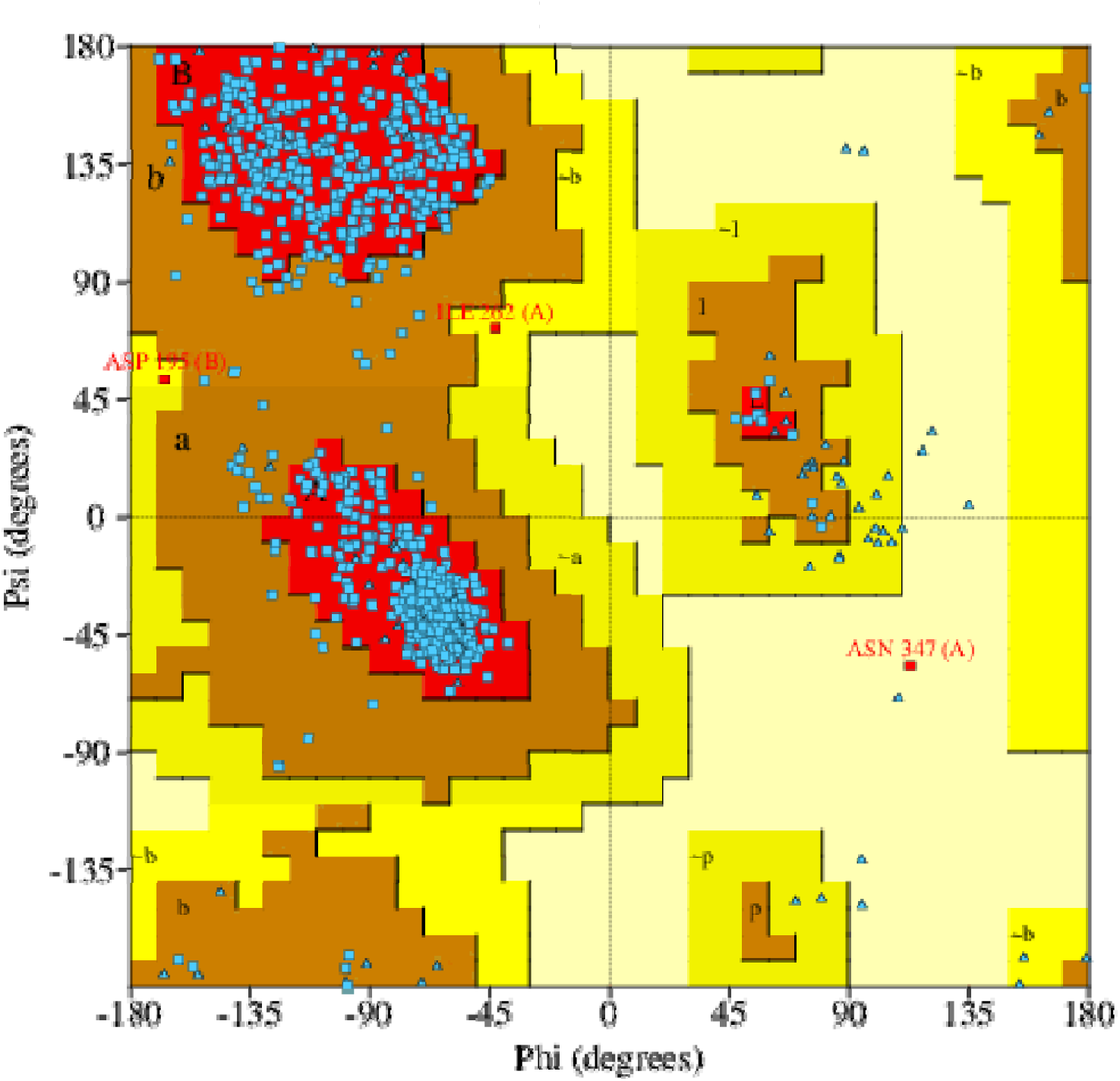
Ramachandran plot for the CNDP2 enzyme structure. Greater than 90% of the residues are located in the most favored regions, which confirms the stereochemical quality of the model. Some residues appear in the allowed outlier regions, including ASN347 and ILE262 in chain A, which are located in flexible loop regions. In contrast, ASP195 in chain B interacts with the catalytic Mn^2+^, which was explicitly considered during docking.

**Figure 3.**
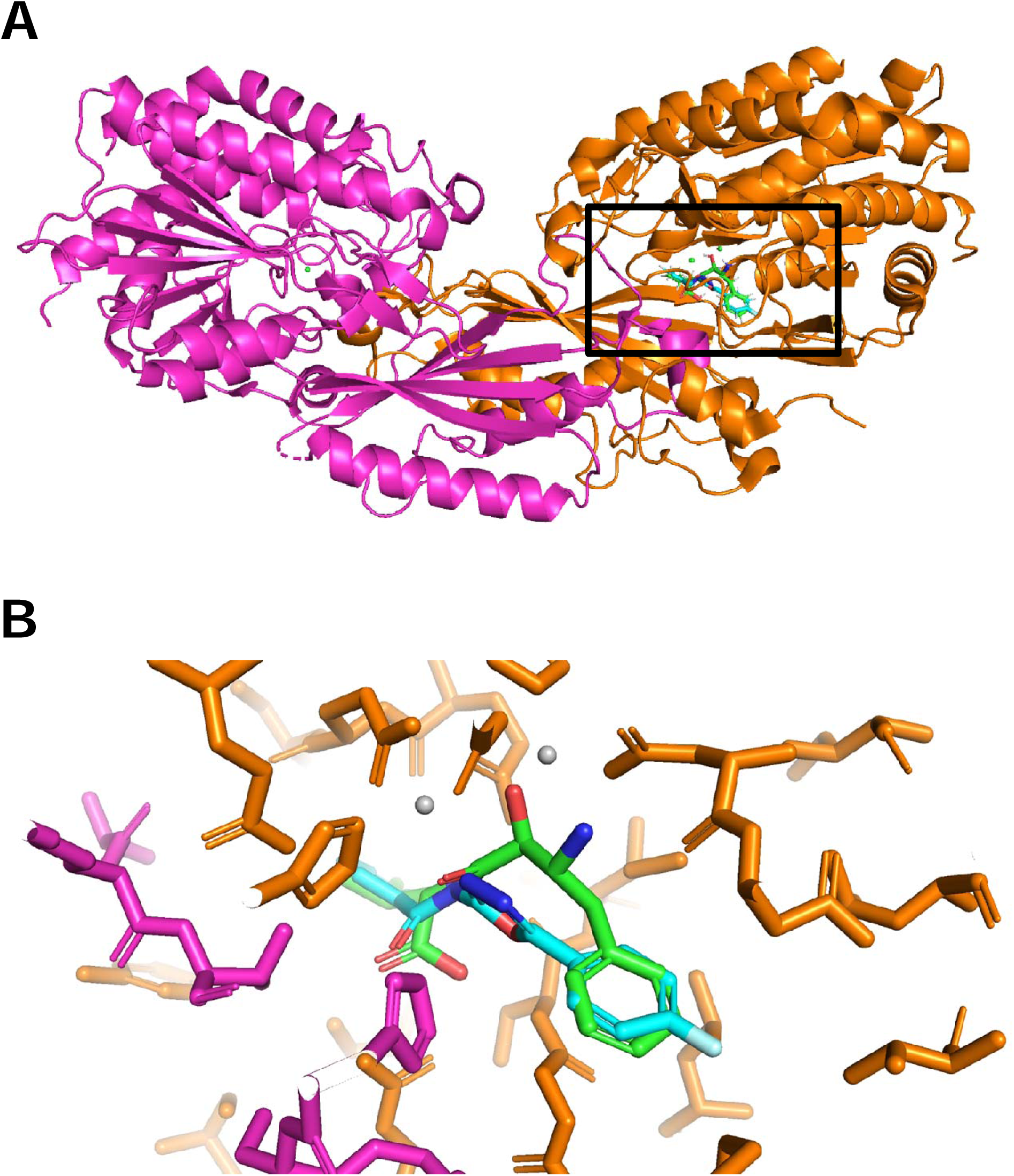
Three-dimensional structure of CNDP2 and orientation of the docked molecules. (A) Overall dimeric structure showing the catalytic cleft. (B) Enlarged view of the black boxed area in (A), which highlights ligand binding within the active pocket. The docked poses for BES (green) and KKL-35 (cyan) substantially overlap within the catalytic pocket.

### Identification of Potential CNDP2 Inhibitors

A structure-based virtual screening of the Selleckchem Bioactive Compound Library-I against CNDP2 revealed two top-ranking compounds: the known inhibitor BES and a novel compound KKL-35, with binding energies of –11.40 kcal/mol and –11.20 kcal/mol, respectively (Table 1). The strong binding affinities of these molecules exceeded those of other screened candidates. BES, which was included as a positive control, exhibited interactions consistent with its reported inhibitory activity. Notably, KKL-35 emerged as a new hit compound and was previously reported to inhibit the trans-translation tagging reaction, thereby resulting in broad-spectrum antibiotic activity [43]. These results suggest that both molecules occupy the catalytic pocket of CNDP2, with KKL-35 representing a novel inhibitor of potential biological relevance.

**Table 1.** Top 10 docking hits against CNDP2 ranked by binding energy scores (kcal/mol).

| Rank | ID | Name | Binding energy score (kcal/mol) |
| --- | --- | --- | --- |
| 1 | drug2490 | Bestatin (BES) | -11.40 |
| 2 | drug2591 | KKL-35 | -11.20 |
| 3 | drug6171 | WAY-660898 | -11.00 |
| 4 | drug5628 | WAY-656494 | -11.00 |
| 5 | drug5970 | WAY-326270-A | -10.90 |
| 6 | drug4125 | IRAK-4 protein kinase inhibitor 2 | -10.90 |
| 7 | drug4843 | 3,6-Dihydroxyflavone | -10.80 |
| 8 | drug5449 | WAY-233765 | -10.70 |
| 9 | drug384 | A-803467 | -10.70 |
| 10 | drug4196 | SIRT7 inhibitor 97491 | -10.60 |

### Binding Interactions and Dynamic Stability of CNDP2–Ligand Complexes

Docking analysis revealed distinct binding modes for BES and KKL-35 (Fig. 4). BES (Fig. 4A) was accommodated within two distinct binding pockets: one formed by His380 and the other by Tyr197 and Gly416 in chain A. Moreover, BES formed multiple hydrogen bonds with Tyr197, Glu166, Ser417, and Arg343 in chain A and Thr330 and His288 in chain B. Direct coordination with the catalytic Mn^2+^ ions was also observed, which is consistent with its known inhibitory mechanism. By contrast, KKL-35 (Fig. 4B) was accommodated within the same Tyr197 and Gly416 pocket and His380 pockets in chain A. It formed hydrogen bonds with Glu166, His380, Arg343, and Ser417 in chain A, and His228 in chain B maintained coordination with an Mn^2+^ ion and formed additional hydrophobic and π-interactions with His380 in chain A and His228 in chain B. These features indicate that both inhibitors targeted the same catalytic site, which may result in enhanced inhibition of CNDP2. To determine the stability of the complexes, 100-ns MD simulations were performed (Fig. 5). RMSD analysis revealed that the protein–ligand complex remained stable throughout the simulation, with values consistently within the 2–3 Å range, which is generally indicative of structural stability in MD studies [44,45]. RMSD analysis indicated that the CNDP2–BES complex was stabilized rapidly and maintained low deviations (∼3 Å), whereas the CNDP2–KKL35 complex exhibited larger fluctuations (∼6–7 Å) after 40 ns (Fig. 5A). The fluctuations indicate that the binding mode of CNDP2-KKL35 changes after 40 ns. RMSF analysis reveals the flexibility of each residue during an MD simulation by quantifying its average positional fluctuations over time [46]. RMSF analysis for a protein–ligand complex provides insights into conformational stability and the local flexibility of the binding-site residues following ligand interaction. RMSF profiles for the CNDP2–ligand complexes revealed less flexibility in the overall structure, including catalytic loop regions, in the presence of KKL-35 compared with BES (Fig. 5B). The radius of gyration (RoG) measures the overall compactness of the protein structure during an MD simulation, and in a protein–ligand complex, it reflects global conformational stability and potential structural changes upon ligand binding [47]. The RoG for the CNDP2–ligand complexes remained compact and steady in the BES system, whereas KKL-35 binding resulted in minor structural looseness (Fig. 5C). Hydrogen bond analysis monitors the number and stability of hydrogen bonds formed during the MD simulation, thus providing insights into key intermolecular interactions that contribute to binding stability [48]. Hydrogen bond analysis confirmed that in the CNDP2–ligand complexes, BES consistently maintained approximately six hydrogen bonds throughout its trajectory, whereas the KKL-35 interactions were fewer and more transient (Fig. 5D). Taken together, these results suggest that BES forms a more stable and persistent inhibitory complex with CNDP2 compared with KKL-35.

**Figure 4.**
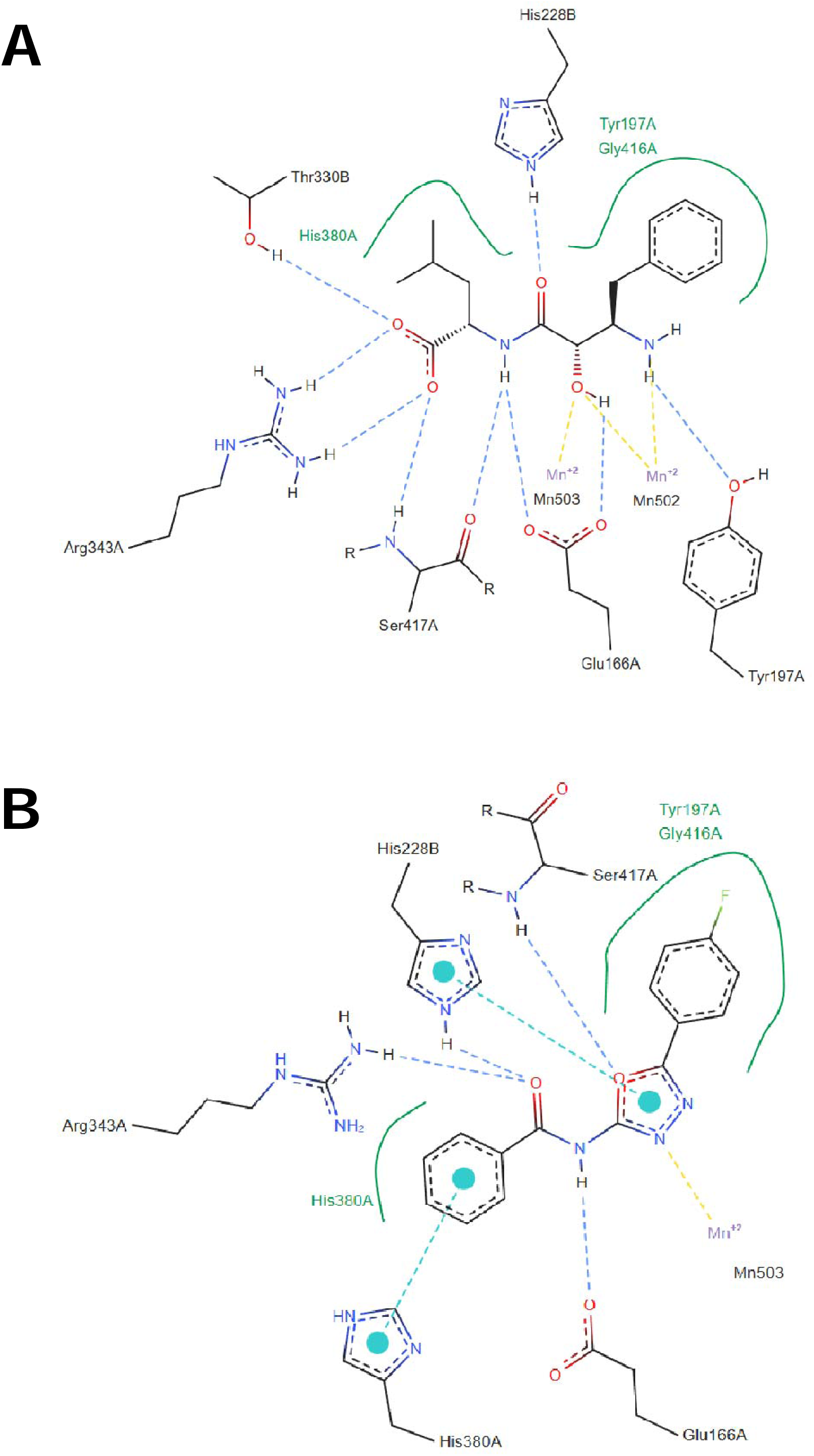
Binding interactions of CNDP2 with docked inhibitors. (A) The BES reference compound. (B) The KKL-35 candidate compound. The green solid lines denote hydrophobic contacts, the yellow dashed lines indicate metal interactions with the catalytic Mn^2+^ ion, the blue dashed lines indicate hydrogen bonds, and the cyan dashed lines with circles represent π–π interactions.

**Figure 5.**
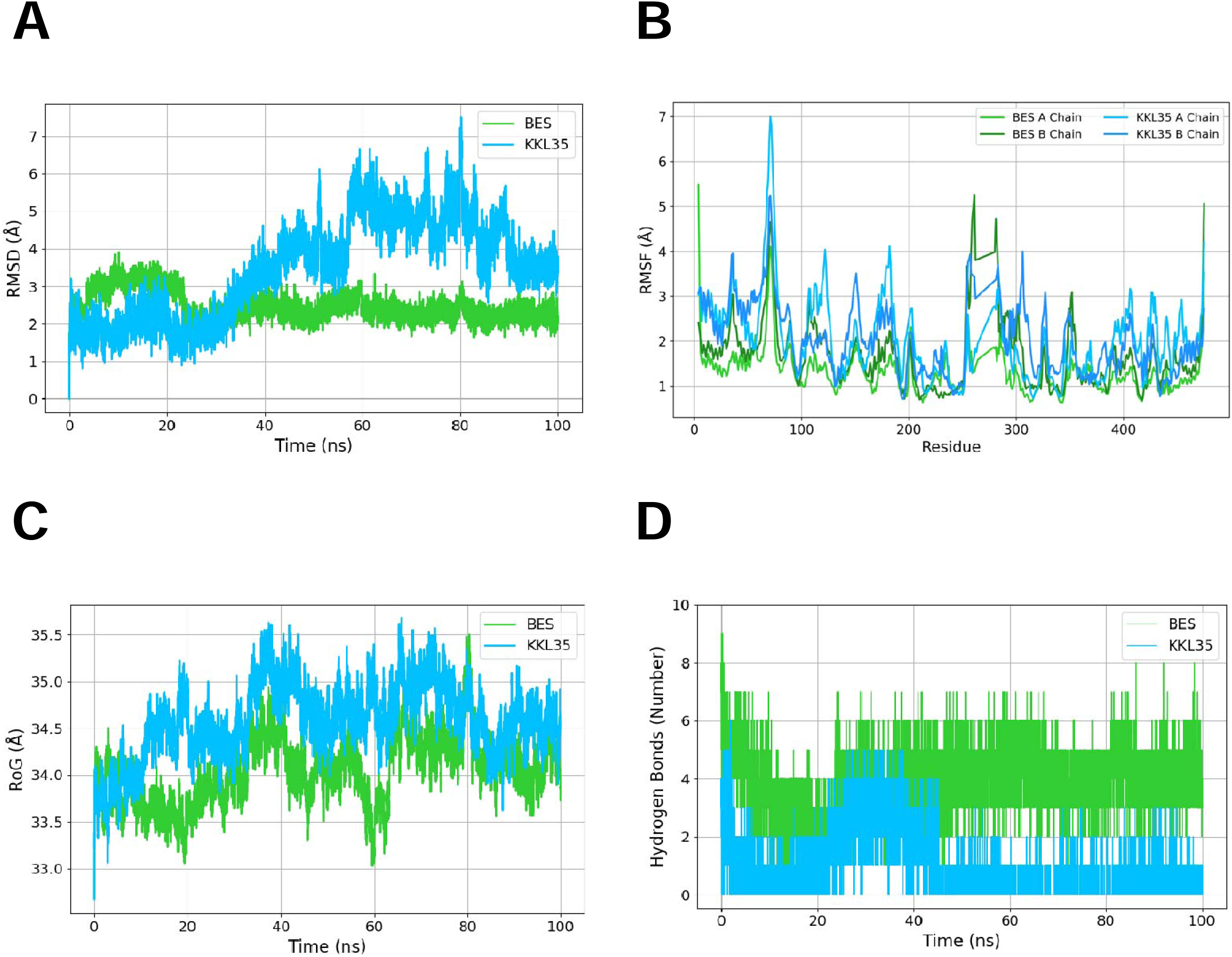
Dynamic behavior of CNDP2–ligand complexes during MD simulations. The yellow–green lines represent the BES complex, and cyan lines denote the KKL-35 complex. (A) Root mean square deviation (RMSD), (B) root mean square fluctuation (RMSF), Lighter lines indicate chain A and darker lines represent chain B. (C) radius of gyration (RoG), and (D) number of hydrogen bonds.

### Binding Free Energy Decomposition and Residue Contributions

To gain further insight into the energetic determinants of ligand recognition, binding free energies were estimated using MM-PBSA calculations, and residue-level energy decomposition was conducted. The Molecular Mechanics Poisson–Boltzmann Surface Area (MM-PBSA) method is used to estimate the binding free energy of biomolecular complexes by combining molecular mechanics energy terms (van der Waals and electrostatic interactions) with solvation free energies, including polar contributions obtained by solving the Poisson–Boltzmann equation and nonpolar contributions derived from the solvent-accessible surface area [49,50]. In addition, to provide an overall estimate of the binding free energy, MM-PBSA allows for residue-level energy decomposition, which highlights individual amino acid contributions to ligand binding. This decomposition analysis is useful for identifying hotspot residues involved in stabilizing the protein–ligand interaction. The total binding free energies (Table 2) revealed that KKL-35 exhibits a weaker binding affinity (–10.05 kcal/mol) compared with BES (–22.59 kcal/mol), which is consistent with the results of docking and MD simulation. In MM-PBSA, the binding free energy (ΔG_bind) is decomposed into several contributions: van der Waals interactions (ΔG_vdW), electrostatic interactions (ΔG_elec), polar solvation energy (ΔG_polar), and nonpolar solvation energy (ΔG_nonpolar). Favorable van der Waals and electrostatic terms represent direct intermolecular interactions between the protein and ligand, whereas polar solvation energy indicates the desolvation penalty required to transfer the ligand from the solvent into the binding pocket. The nonpolar solvation term, which is estimated from the solvent-accessible surface area, typically favors binding because of hydrophobic effects. The results indicated that KKL-35 and BES were stabilized by van der Waals and electrostatic interactions; however, BES exhibited a much stronger electrostatic contribution (–92.73 kcal/mol) compared with that of KKL-35 (–29.32 kcal/mol), which was partially offset by the unfavorable polar solvation energy (+109.15 kcal/mol). In contrast, KKL-35 exhibited weaker electrostatic stabilization, but incurred a smaller desolvation penalty, which ultimately resulted in a less favorable ΔG_bind compared with BES.

**Table 2.** MM-PBSA binding free energy decomposition of CNDP2–ligand complexes.

| Compound | $\Delta G$ binding | $\Delta G$ | $\Delta G$ bind van | $\Delta G$ bind | $\Delta G$ polar | $\Delta G$ nonpolar | $\Delta G$ solvation |
| --- | --- | --- | --- | --- | --- | --- | --- |
|  |  | electrostatic | der Waals | gas phase | solvation | solvation |  |
| KKL-35 | -10.05 | -29.32 | -43.62 | -72.94 | 66.41 | -3.52 | 62.89 |
| BES | -22.59 | -92.73 | -35.01 | -127.75 | 109.15 | -3.99 | 105.16 |

Decomposition of the MM-PBSA terms indicated that van der Waals and electrostatic interactions favor complex formation for both ligands, whereas the polar solvation energy opposes binding, with the nonpolar solvation term providing a modest favorable contribution [51]. The residue-level energy decomposition (Fig. 6) revealed a set of shared stabilizing contacts (ΔG ≤ –0.5 kcal mol^-1^), including GLY169, GLY415, SER417, ILE213, and ILE418 in chain A. In contrast, KKL-35 exhibited unfavorable contributions from GLU167 in chain A (+4.18 kcal mol^-1^) and small penalties at GLU414 (+0.24 kcal mol^-1^) and HIS445 (+0.37 kcal mol^-1^) in chain A. Notably, the two catalytic Mn^2+^ ions did not uniformly stabilize the complexes. For BES, Mn502 was strongly unfavorable (+11.14 kcal mol^-1^), whereas Mn503 was stabilizing (–6.81 kcal mol^-1^). The pattern was reversed for KKL-35, in which Mn502 was stabilizing (–8.27 kcal mol^-1^) and Mn503 was destabilizing (+3.99 kcal mol^-1^). These opposing signs indicate distinct metal-coordination/solvation environments for the two ligands, rather than a generic metal-assisted stabilization. Taken together, BES achieved a more favorable ΔG_bind by combining stronger electrostatics with the favorable engagement of one Mn site, whereas KKL-35 suffers from penalties at GLU167 and at one of the Mn centers, which is consistent with its overall weaker affinity.

**Figure 6.**
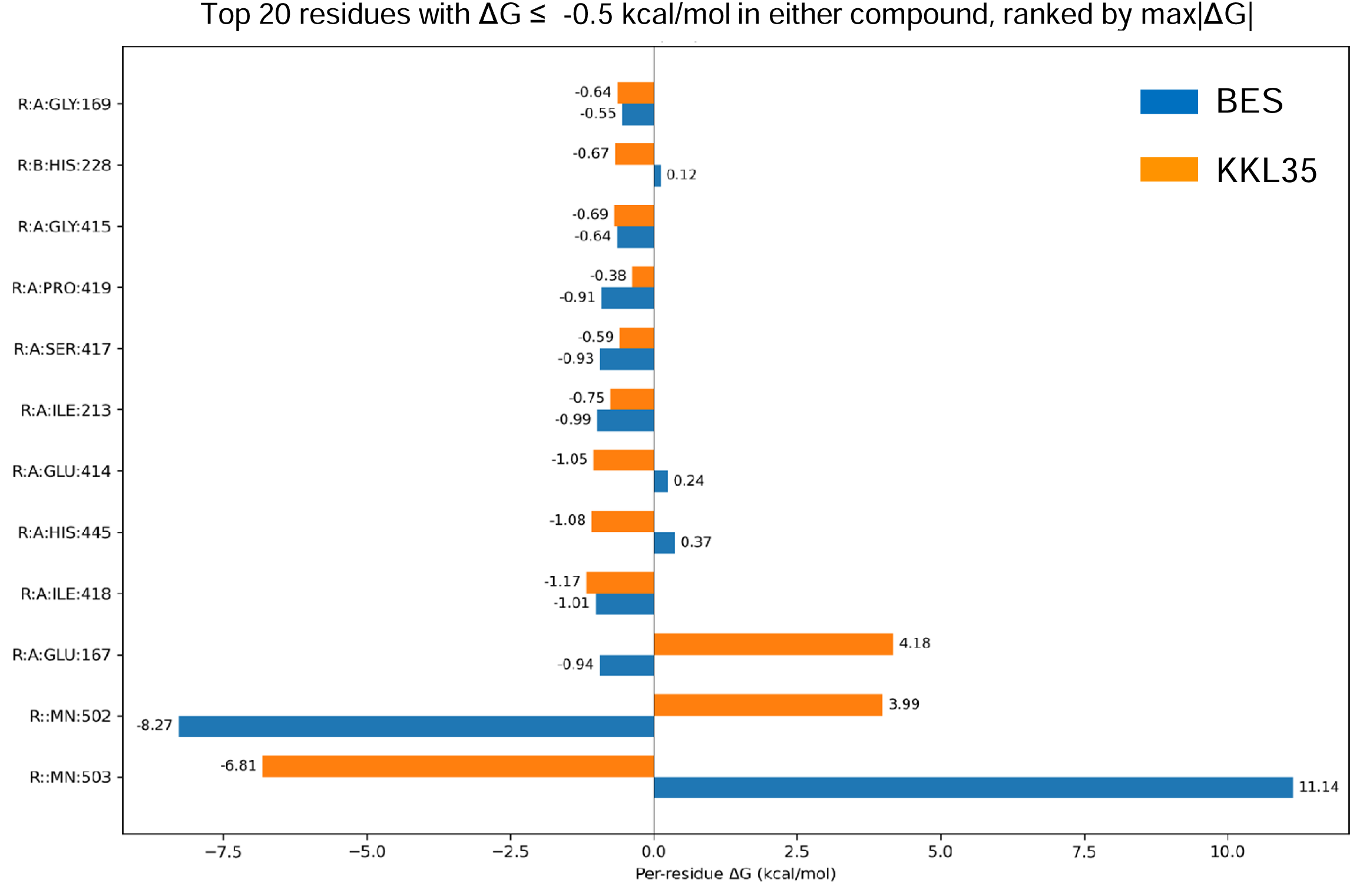
Per-residue binding free energy decomposition analysis of the CNDP2–ligand complexes, which highlight key amino acid contributions to the stability of BES and KKL-35 binding. The vertical axis indicates amino acid residues and catalytic Mn^2+^ ions, whereas the horizontal axis shows the per-residue binding free energy contribution (ΔG, kcal/mol). The analysis highlights key residues that contribute to the stability of BES (blue) and KKL-35 (orange) binding. The top 20 residues are displayed with ΔG ≤ –0.5 kcal/mol in either compound, ranked by the maximum absolute contribution (max|ΔG|).

### ADMET and Drug-Likeness Analysis

The ADMET and drug-likeness properties of BES and KKL-35 were determined using SwissADME (Table 3). The two compounds satisfied major drug-likeness filters (Lipinski, Ghose, Veber, Egan, and Muegge rules) without violations and achieved a bioavailability score of 0.55 [23,52–55]. Neither triggered PAINS or Brenk alerts, which confirmed the absence of problematic substructures [24,56]. KKL-35 had a favorable synthetic accessibility score (2.64) compared with BES (3.10), which suggests that KKL-35 may be more amenable to chemical optimization during lead development [54]. The physicochemical parameters, including molecular weight, topological polar surface area (TPSA), and rotatable bond count, are important determinants of oral drug-likeness. KKL-35, with a molecular weight of 317.7 Da, a TPSA of 68.02 Å², and four rotatable bonds, was within the range associated with good passive permeability and oral bioavailability [23,53,57]. In contrast, BES showed a higher polarity (TPSA 112.65 Å²) and a greater degree of conformational flexibility (nine rotatable bonds), which may limit its membrane permeability and reduce binding affinity. Lipophilicity is expressed as consensus Log P and reflects the balance between hydrophobic and hydrophilic characteristics, which strongly affects absorption, distribution, and binding to plasma proteins. KKL-35 exhibited moderate and well-balanced lipophilicity (3.44), which favors solubility and permeability, whereas BES showed lower lipophilicity (0.79). Although beneficial for solubility, the latter may compromise passive diffusion across lipid membranes. Solubility predictions further highlight these differences: BES was predicted to be highly soluble, which facilitates formulation and dissolution, whereas KKL-35 was only moderately soluble, indicating a potential challenge that may require formulation strategies, such as salt formation or nanoparticle delivery systems. Pharmacokinetic predictions suggested high gastrointestinal absorption for both compounds, which is consistent with their overall compliance with the drug-likeness rules; however, blood–brain barrier (BBB) penetration diverged notably. KKL-35 was predicted to cross the BBB, indicating its potential for central nervous system (CNS) exposure, whereas BES was not, which may reduce CNS-related off-target effects, but also limit the efficacy for CNS targets. In addition, transporter interactions supported favorable pharmacokinetics. Neither compound was predicted to be a substrate for P-glycoprotein (P-gp), an efflux pump that reduces oral bioavailability and brain penetration. Cytochrome P450 (CYP) inhibition profiles provided insights into its drug–drug interaction potential. KKL-35 was predicted to inhibit CYP1A2 and CYP2C19 activities, which raised the possibility of metabolic interactions with co-administered drugs metabolized by these isoforms. By contrast, BES showed no predicted inhibition among the major CYP450 enzymes, suggesting a lower liability for pharmacokinetic drug–drug interactions. The high lipophilicity of KKL-35 likely contributes to its susceptibility to hepatic CYP-mediated metabolism, consistent with the predicted inhibition of CYP1A2/2C19 [58].

**Table 3.** Physicochemical, pharmacokinetic, and drug-likeness profiles of KKL-35 and BES.

|  | Compound | KKL-35 | BES |
| --- | --- | --- | --- |
| Physicochemical properties | Formula | C15H9ClFN3O2 | C16H24N2O4 |
|  | Molecular weight | 317.7 | 308.37 |
|  | #Heavy atoms | 22 | 22 |
|  | #Aromatic heavy atoms | 17 | 6 |
|  | Fraction Csp3 | 0 | 0.5 |
|  | #Rotatable bonds | 4 | 9 |
|  | #H-bond acceptors | 5 | 5 |
|  | #H-bond donors | 1 | 4 |
|  | Molar refractivity | 78.92 | 83.31 |
|  | TPSA (Å<sup>2</sup>) | 68.02 | 112.65 |
| Lipophilicity | Consensus Log P | 3.44 | 0.79 |
|  | Water solubility | Moderately soluble | Very soluble |
| Pharmacokinetics | GI absorption | High | High |
|  | BBB permeant | Yes | No |
|  | P-gp substrate | No | No |
|  | CYP1A2 inhibitor | Yes | No |
|  | CYP2C19 inhibitor | Yes | No |
|  | CYP2C9 inhibitor | No | No |
|  | CYP2D6 inhibitor | No | No |
|  | CYP3A4 inhibitor | No | No |
|  | log Kp (cm/s) | -5.89 | -8.86 |
| Druglikeness | Lipinski | Yes; 0 violation | Yes; 0 violation |
|  | Ghose | Yes | Yes |
|  | Veber | Yes | Yes |
|  | Egan | Yes | Yes |
|  | Muegge | Yes | Yes |
|  | Bioavailability Score | 0.55 | 0.55 |
| Medicinal chemistry | PAINS #alerts | 0 | 0 |
|  | Brenk #alerts | 0 | 0 |
|  | Synthetic Accessibility | 2.64 | 3.1 |

Skin permeability (log Kp) values, which reflect compound diffusion across biological barriers, were more favorable for KKL-35 (–5.89 cm/s) compared with BES (–8.86 cm/s), which is consistent with its lower polarity and superior lipophilicity profile [59]. Taken together, these results indicate that while both molecules exhibit acceptable drug-likeness and oral bioavailability potential, KKL-35 exhibits superior balance in terms of permeability, lipophilicity, and synthetic accessibility, whereas BES offers higher aqueous solubility and a reduced risk of CYP-mediated drug–drug interactions.

The SwissTargetPrediction report presents a pie chart illustrating the distribution of predicted target classes for BES as the query compound (Supplementary Fig. 1). The analysis indicated that BES is a potential ligand for major proteases (66.7%, 10 of the top 15 predicted targets). We also identified aminopeptidases, including aminopeptidase N, aminopeptidase B, and leucine aminopeptidase, as well as related enzymes, such as matrix metalloproteinase-2 and leukotriene A4 hydrolase, which are among the top five predicted targets for BES, and predicted with high probability (Table 4). By contrast, the SwissTargetPrediction report for KKL-35 indicates that proteases are not a prominent class of targets and no specific high-probability target could be identified (Table 5).

**Table 4.** Protease-related molecular targets of BES predicted by SwissTargetPrediction.

| Target | Common name | Probability |
| --- | --- | --- |
| Aminopeptidase N | ANPEP | 0.952247188 |
| Aminopeptidase B (by homology) | RNPEP | 0.952247188 |
| Matrix metalloproteinase 2 | MMP2 | 0.952247188 |
| Leucine aminopeptidase | LAP3 | 0.952247188 |
| Leukotriene A4 hydrolase | LTA4H | 0.952247188 |
| Aspartyl aminopeptidase | DNPEP | 0.135202128 |
| Angiotensin-converting enzyme | ACE | 0.119403562 |
| Dipeptidyl peptidase IV | DPP4 | 0.119403562 |
| Xaa-Pro dipeptidase | PEPD | 0.111501865 |
| Calpain 1 | CAPN1 | 0.111501865 |
| Neprilysin (by homology) | MME | 0.111501865 |
| Beta-secretase 1 | BACE1 | 0.111501865 |
| Dipeptidyl peptidase VIII | DPP8 | 0.111501865 |
| Dipeptidyl peptidase IX | DPP9 | 0.111501865 |
| Methionine aminopeptidase 2 | METAP2 | 0.111501865 |
| Carboxypeptidase B | CPB1 | 0.111501865 |
| Renin | REN | 0.111501865 |
| Xaa-Pro aminopeptidase 2 | XPNPEP2 | 0.111501865 |
| Prolyl endopeptidase | PREP | 0.111501865 |
| Fibroblast activation protein alpha | FAP | 0.111501865 |
| Renal dipeptidase | DPEP1 | 0.111501865 |
| Beta secretase 2 | BACE2 | 0.111501865 |
| Caspase-1 | CASP1 | 0.111501865 |

**Table 5.** Protease-related molecular targets of KKL-35 predicted by SwissTargetPrediction.

| Target | Common name | Probability |
| --- | --- | --- |
| Thrombin and coagulation factor X | F10 | 0.111501865 |
| Caspase-3 | CASP3 | 0.111501865 |
| Matrix metalloproteinase 13 | MMP13 | 0.111501865 |
| Gamma-secretase | PSEN2 | 0.111501865 |
| Cathepsin (V and K) | CTSV | 0.111501865 |
| Cathepsin L | CTSL | 0.111501865 |
| Cathepsin G | CTSG | 0.111501865 |
| Prolyl endopeptidase | PREP | 0.111501865 |
| Fibroblast activation protein alpha (by homology) | FAP | 0.111501865 |
| Caspase-7 | CASP7 | 0.111501865 |

### *In Vitro* Inhibitory Analysis of Ligands

To determine whether KKL-35 suppresses CNDP2 enzymatic activity, we evaluated its effect on the hydrolysis of Cys-Gly into Cys and Gly using purified CNDP2. Cys was derivatized with NEM and quantified as Cys-NEM. KKL-35 inhibited CNDP2 activity in a dose-dependent manner, yielding IC_50_ values of 118 μM for Cys-NEM and 96 μM for glycine, thus confirming its potential as a CNDP2 inhibitor (Fig. 7A). In contrast, the established inhibitor BES was much more potent, with IC_50_ values of 1.0 μM and 1.2 μM (Fig. 7B). These results are consistent with those of our MD simulations, which predicted markedly stronger binding of BES to CNDP2 compared with KKL-35. Taken together, these results confirm that although less potent than BES, KKL-35 represents an effective CNDP2 inhibitor.

**Figure 7.**
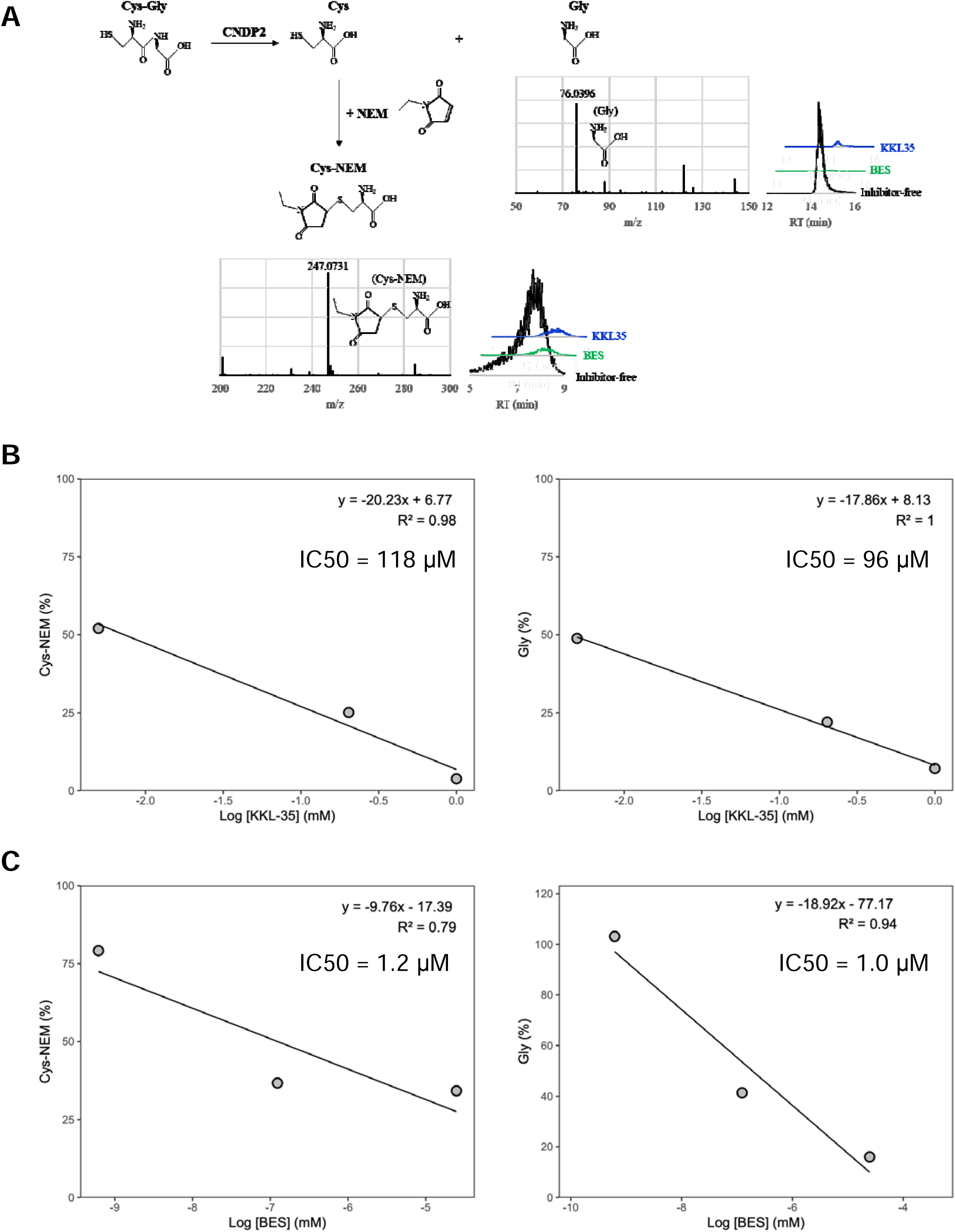
LC–MS analysis of CNDP2-mediated Cys-Gly hydrolysis and its inhibition by BES and KKL-35. (A) LC–MS analysis of CNDP2-catalyzed hydrolysis of Cys-Gly and its inhibition by BES and KKL-35. Cys-Gly was hydrolyzed by CNDP2 to produce cysteine (Cys) and glycine (Gly). Cys was derivatized with *N*-ethylmaleimide (NEM) to form Cys–NEM. Mass spectra confirmed the identities of Gly (*m/z* 76.0396) and Cys–NEM (*m/z* 247.0731). Extracted ion chromatograms showing reduced levels of Gly and Cys–NEM in the presence of BES or KKL-35 compared with the inhibitor-free control. (B, C) Inhibition of CNDP2 enzymatic activity by KKL-35 (B) and BES (C). Dose-dependent inhibition of Cys-Gly hydrolysis was assessed by measuring the production of Cys (measured as Cys-NEM after derivatization) and Gly. KKL-35 exhibited IC_50_ values of 118 μM for Cys-NEM and 96 μM for Gly, whereas BES showed markedly higher potency, with IC_50_ values of 1.2 μM and 1.0 μM, respectively.

## 4. Discussion

Carnosine dipeptidase 2 (CNDP2) represents an attractive therapeutic target because of its roles in maintaining intracellular cysteine availability for glutathione synthesis and supporting tumor cell proliferation through nutrient recycling. Therefore, inhibiting CNDP2 may disrupt the redox balance and sensitize tumor cells to oxidative stress, which is a rational strategy for cancer therapy; however, the established inhibitor BES is limited by poor selectivity and suboptimal drug-like properties. BES is a broad-spectrum peptidase inhibitor that targets multiple enzymes, which raises concerns regarding the off-target toxicity [13–15]. Moreover, its high TPSA and numerous rotatable bonds compromise membrane permeability and increase the entropic costs of binding (Table 3), resulting in a drug-likeness profile that is not ideal for development.

Using an *in silico* screen, we identified KKL-35 as a novel CNDP2-binding molecule. Computational analyses suggested that KKL-35 engages in distinct Mn^2+^ coordination and favorable hydrophobic and hydrogen-bond interactions within the catalytic pocket (Fig. 4). MD simulation and MM-PBSA analysis indicated that the CNDP2–KKL-35 complex has less stability compared with the CNDP2–BES complex (Fig. 5 and Table 2), which is consistent with the inhibitory activity observed in enzymatic assays (Fig. 7). Although the potency of KKL-35 was lower compared with that of BES, KKL-35 functions as a bona fide CNDP2 inhibitor, thus supporting the predictive power of the computational workflow and providing an entry point for chemical optimization.

Despite its weaker intrinsic activity, KKL-35 holds considerable advantages as a potential lead scaffold. It has more favorable physicochemical properties than BES, including a lower TPSA, fewer rotatable bonds, and balanced lipophilicity (Table 3), all of which support improved oral bioavailability and membrane permeability. Its synthetic accessibility score further suggests that structural optimization is feasible (Table 3). KKL-35 was predicted to cross the BBB (Table 3). Although not essential for peripheral cancers, this broadens its potential applications to tumors of the CNS as well as neurodegenerative disorders, in which glutathione metabolism and oxidative stress play central roles in pathology. Conversely, unintended brain exposure in diseases outside of the nervous system may pose safety risks, which highlights the importance of tailoring permeability during future optimization.

Residue-level energy decomposition revealed hotspot residues, including Tyr197, His228, His380, Arg343, and Glu166, which markedly contribute to ligand stabilization (Fig. 6). These insights provide a rational roadmap for enhancing KKL-35 activity through medicinal chemistry, such as reinforcing hydrogen bonding and π–π interactions with these residues. Although BES benefits from stronger binding affinity, its broad inhibitory profile and unfavorable physicochemical characteristics limit its therapeutic applicability. In contrast, KKL-35 combines the verifiable inhibitory activity with superior drug-like features, making it a promising starting point for lead optimization.

In conclusion, BES is a valuable reference compound, but is unsuitable as a therapeutic scaffold because of its nonselectivity and poor drug-likeness. KKL-35, although less potent, represents a validated CNDP2 inhibitor with much room for improvement. Its favorable ADMET properties and tractability for chemical modification position it as a lead candidate for the development of next-generation CNDP2 inhibitors. Therefore, the present study establishes KKL-35 not as a final solution, but as a lead compound for rational optimization toward potent and selective CNDP2-targeted therapeutics.

## Declaration of competing interest

The authors declare no conflicts of interest.

## Notes

### Competing Interest Statement

The authors have declared no competing interest.

