## Supplementary Fig 1 for "Identification of KKL-35 as a novel carnosine dipeptidase 2 (CNDP2) inhibitor by *in silico* screening"

### Slide 1
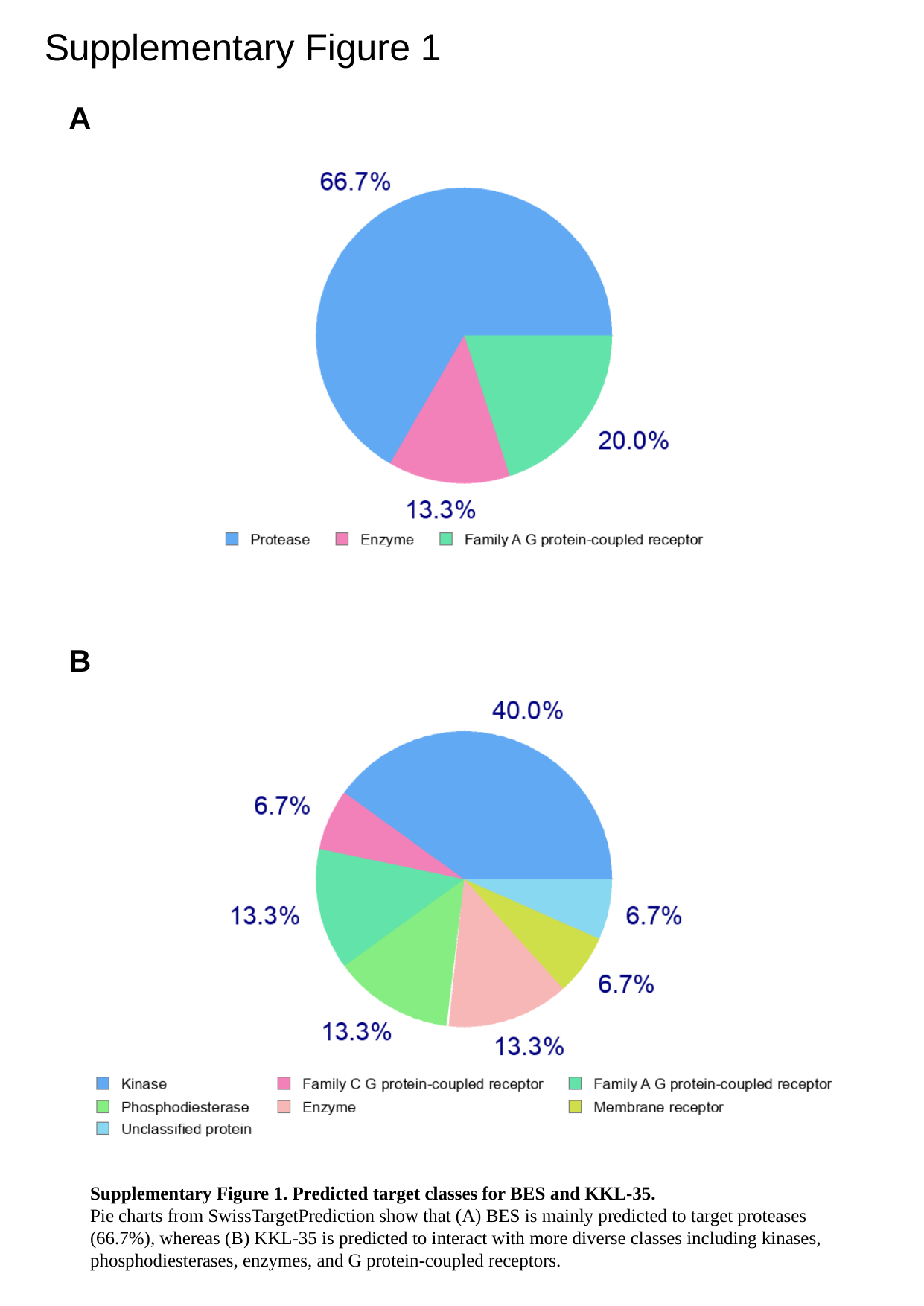

Supplementary Figure 1
A
B
Supplementary Figure 1. Predicted target classes for BES and KKL-35.
Pie charts from SwissTargetPrediction show that (A) BES is mainly predicted to target proteases (66.7%), whereas (B) KKL-35 is predicted to interact with more diverse classes including kinases, phosphodiesterases, enzymes, and G protein-coupled receptors.
